# Interspecies Killing Activity of *Pseudomonas syringae* Tailocins

**DOI:** 10.1101/2022.07.29.502058

**Authors:** Savannah L. Weaver, Libin Zhu, Sadhana Ravishankar, Meara Clark, David A. Baltrus

## Abstract

Tailocins are ribosomally synthesized bacteriocins, encoded by bacterial genomes, but originally derived from bacteriophage tails. As with both bacteriocins and phage, tailocins are largely thought to be species-specific with killing activity often assumed to be directed against closely-related strains. Previous investigations into interactions between tailocin host range and sensitivity across phylogenetically diverse isolates of the phytopathogen *Pseudomonas syringae* have demonstrated that many strains possess intraspecific tailocin activity and that this activity is highly precise and specific against subsets of strains. However, here we demonstrate that at least one strain of *P. syringae*, USA011R, defies both expectations and current overarching dogma because tailocins from this strain possess broad killing activity against other agriculturally significant phytopathogens such as *Erwinia amylovora* and *Xanthomonas perforans* as well as against the clinical human pathogen *Salmonella choleraesuis*. Moreover, we show that the full spectrum of this interspecific killing activity is not conserved across closely related strains with data suggesting that even if tailocins can target different species, they do so with different efficiencies. Our results reported herein highlight the potential for and phenotypic divergence of interspecific killing activity of *P. syringae* tailocins and establish a platform for further investigations into the evolution of tailocin host range and strain specificity.

## Introduction

Broad-spectrum antibiotics have revolutionized the treatment of clinical and veterinary diseases since their development in the 19^th^ century. However, due to realized worries about selecting for the evolution and dissemination of resistance genes across environments, similar antibiotics have been used sparingly to prevent or treat infections by phytopathogens in plant-based agricultural settings (McManus et al. 2002; Stockwell and Duffy 2012). Furthermore, as important roles of natural microbiomes in fostering healthy physiology and development have been uncovered across clinical and agricultural systems, there are increased fears of detrimental off-target effects of the widespread use of broad-spectrum antibiotics (Andersson and Hughes 2014; Aslam et al. 2021). Lastly, in contrast to their use for the treatment of human and animal infections, that application of antibiotics is often not effective for treatment once diseased crops are seen under field conditions (Stockwell and Duffy 2012). Taken together, what is needed is the development of programmable strain-specific antimicrobial compounds that can be effectively applied as prophylactic treatments to prevent foodborne pathogens from entering the food chain through produce.

Numerous potential avenues are being developed as effective prophylactic treatments to prevent phytopathogen infection in agriculture. For instance, phage therapy has been successfully used to treat multidrug-resistant bacterial infections in humans and shows great potential in plants as well (Buttimer et al. 2017). However, exposure to environmental stressors (i.e. soil pH, UV radiation) could lead to inactivation of phage and lack of killing efficiency under environmental conditions due to challenges with genome replication (Buttimer et al. 2017). Hesitation also exists, whether the worries are realistic or not, about unpredictable future evolutionary dynamics and the uncontrolled nature of phage release into the environment (Meaden and Koskella 2013). Alternatively, bacteriocins are natural antimicrobials produced by bacteria that are largely thought to precisely target and kill strains of the same or closely related species (Riley and Wertz 2002). Since they are proteinaceous and non-replicative, it is possible that they could be applied as treatments under agricultural conditions with a reduced risk of environmental stress interfering with the effectiveness and with reduced worries over evolutionary dynamics (Patz et al. 2019; Cotter, Ross, and Hill 2013; Baltrus et al. 2022; Sheth et al. 2016). Bacteriocins have also proven effective in the prevention or treatment for various plant diseases, but their use is somewhat limited by the necessity to find and characterize compounds that are effective against bacterial strains of interest (e.g. Rooney et al. 2020). Further development of a bacteriocin based platform where molecules could be engineered or programmed with designer killing spectra would be an exceptional next step towards their widespread use as prophylactic compounds against a variety of agricultural pathogens.

Phage tail-derived bacteriocins, hereafter referred to as tailocins, combine the specificity and effectiveness of killing of both phage and bacteriocins and display great potential as the basis of a programmable bacteriocin platform (Patz et al. 2019; Ghequire and De Mot 2015). Expression of tailocin genes is triggered by DNA damage, with tailocins thought to be used to outcompete strains for the same niche space (Gillor, Etzion, and Riley 2008; Riley and Wertz 2002).Tailocins are assembled inside producer cells and are released via cell lysis, bind to receptors on target cells (for Gram negative targets, receptors are often the lipopolysaccharide (LPS), and use a contractile sheath to puncture cell membranes and disrupting cellular integrity, thereby killing the target cell by lysis (Patz et al. 2019; Ghequire and De Mot 2015). It is currently thought that, so long as it can sufficiently bind to a target cell, that a single tailocin molecule is sufficient to eliminate a single bacteria cell, although see (Kandel, Baltrus, and Hockett 2020). This “one hit, one kill” mechanism provides an efficient and precise means to eliminate target cells without necessarily having to worry about minimum inhibitory concentrations in the context of effectiveness (Patz et al. 2019). Lastly, we have recently demonstrated that prophylactic treatment of plants with tailocins is effective at protecting *Nicotiana benthamiana* from infection by *Pseudomonas syringae* pv. *syringae* B728A (Baltrus et al. 2022), which mirrors results from *P. fluorescens* in controlling bacterial-spot disease in tomatoes (Príncipe et al. 2018).

To use tailocins as programmable antimicrobials it is important that we better understand how host range is determined and how it can change (Cotter, Ross, and Hill 2013; Riley and Wertz 2002). To this point, investigations of killing activity have shown the targeting range for tailocins is potentially broader than previously imagined. One study showed the killing range of the phage tail-like bacteriocin isolated from *Burkholderia cenocepacia* to be quite broad, including killing activity against *Pseudomonas aeruginosa* (Yao et al. 2017) and R-type pyocins from *Pseudomonas aeruginosa* have been shown to target both *Neisseria gonorrhoeae* and *Neisseria meningitidis* (Blackwell and Law 1981). Although the tailocins of *P. syringae* and *P. fluorescens* arose from independent evolutionary events compared to those of *Burkholderia* and *P. aeruginosa*, data from Principe et al. also suggest that these molecules could maintain interspecific activity against a *Xanthomonas* strain (Príncipe et al. 2018).

Previous characterization of tailocins produced by the phytopathogen *Pseudomonas syringae* and have shown that these tailocins can be effectively used as a prophylactic treatment for plant disease (Baltrus et al. 2022). Previous research suggested that the intraspecific killing spectrum of these tailocins is specific across closely related strains within the *P. syringae* complex (Baltrus et al. 2018), but here we expand on these results and report that the killing activity of tailocins produced by *P. syringae* strain USA011R can be interspecific. Tailocins from this strain can kill phytopathogenic *Erwinia* and *Xanthomonas*, and even demonstrate specific activity against a single *Salmonella* species. These results bring us closer to understanding the mechanisms behind tailocin killing spectrum evolution and highlight how closely related strains may possess subtle to dramatic phenotypic differences in interspecific killing. Identifying the determinants of host range changes at the molecular level could enable precise engineering of tailocin molecules with designer killing specificities.

## Results

### Cross-species Killing Activity of *Pseudomonas syringae* Tailocins

We tested the killing activity of *P. syringae* USA011R tailocins by extracting tailocin molecules and performing soft agar overlay assays against *P. syringae* pv. *syringae* B728a, *Erwinia amylovora* ATCC 29780, *Xanthomonas perforans* DBL942, and *Salmonella enterica* subsp. *enterica* serovar *choleraesuis* strain ETS 34 (ATCC 10708) (Fig. 1). A previous study developed strain USA011R as a platform for investigating the genetic basis of tailocin targeting, and demonstrated that tailocin Receptor Binding Protein (hereafter Rbp) and chaperone are necessary for killing of target strains and for determining the host range of tailocins (Baltrus et al. 2018, 2022). In a previous publication, we created deletion strains whereby tailocin killing activity was eliminated when the Rbp and chaperone genes were deleted from USA011R, but also complemented this deletion *in cis* through the reintroduction of these two loci (Baltrus et al. 2022). We use these strains again for the data presented in Figure 1, which shows that the tailocin from USA011R can target and kill a broader array of target strains than previously thought including strains within the genera *Erwinia, Xanthomonas*, and *Salmonella*. This broader activity is eliminated in the Rbp/chaperone mutant, and restored in the complementation strain. Therefore, the Rbp and chaperone of strain USA011R are both necessary for tailocin killing of *P. syringae* strains as well as for the interspecific killing activity shown in Figure 1.

**Figure 1:**
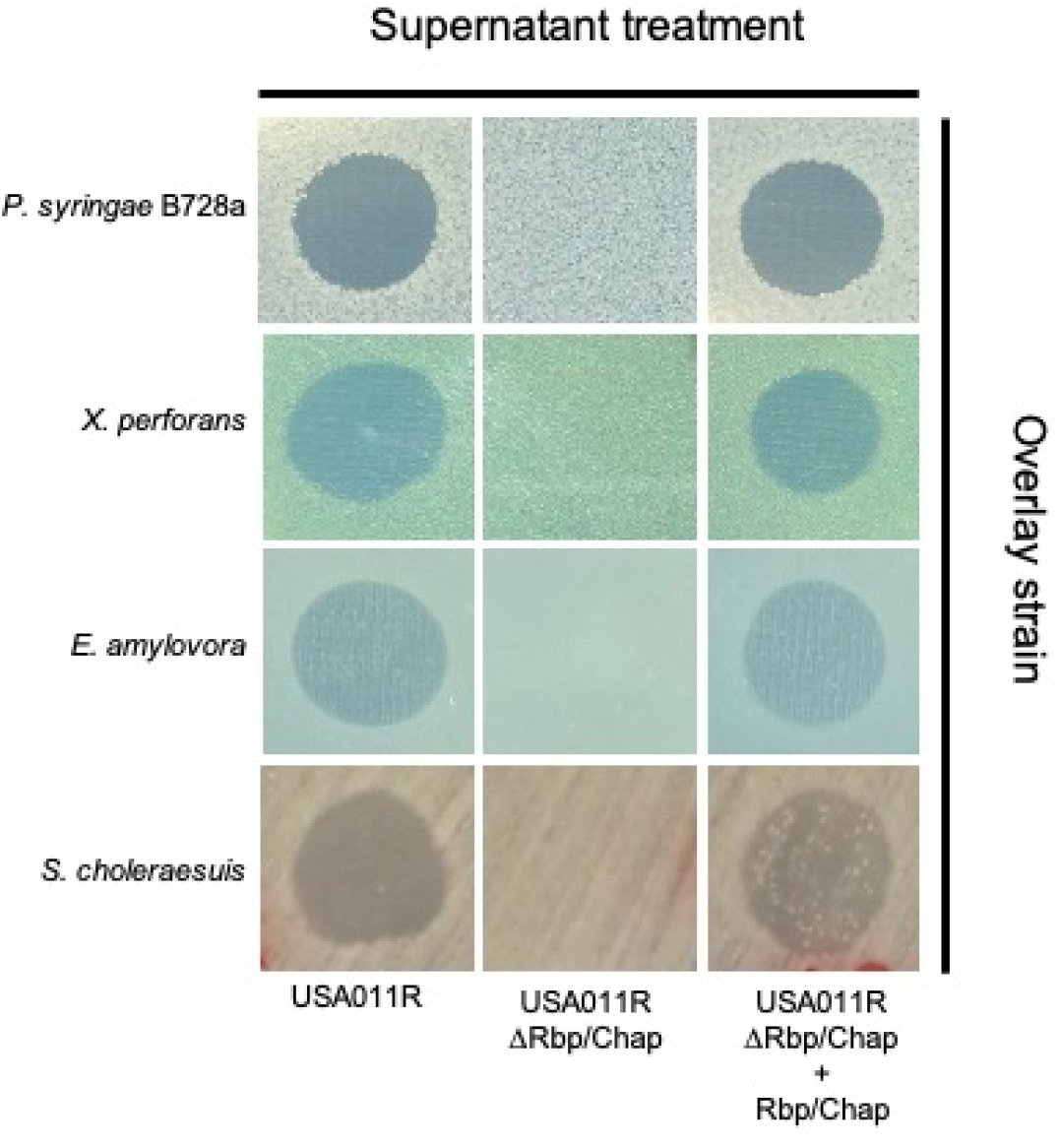
*Pseudomonas syringae* strain USA011R Tailocins Display Interspecies Killing Activity. Overlay experiments are shown as per Hockett and Baltrus (2017) and Baltrus et al. (2022). Killing activity is indicated by a clearing in the lawn of target bacteria. Supernatant treatments from three different strains are shown along the horizontal axis: USA011R, an Rbp/chaperone deletion mutant labelled USA011RΔRbp/Chap, and a complemented strain of this deletion mutant labelled USA011RΔRbp/Chap+Rbp/Chap. Target strains of these assays are represented on the vertical axis. Data is representative of at least three independent overlay experiments. Pictures used for this figure were modified to enhance contrast, but original pictures can be found as supplemental figure S1A, S1B, S1C, and S1D.

To further vet killing activity of strain USA011R, we tested for killing or inhibitory activity of this strain against a variety of other clinically and agriculturally relevant human pathogen strains (Table 1). Surprisingly, we found inhibitory activity of tailocins from USA011R only against *S. choleraesuis* and not against any other foodborne pathogens tested including *Salmonella, E. coli* O157:H7, or *Listeria monocytogenes* strain. Therefore, while the interspecific killing spectrum of tailocins from USA011R is broader than previously imagined, this activity parallels what has been demonstrated for tailocins within a species (Baltrus et al. 2018) with targeting likely occurring at the level of individual strains rather against all strains within a genus or species.

**Table 1:**
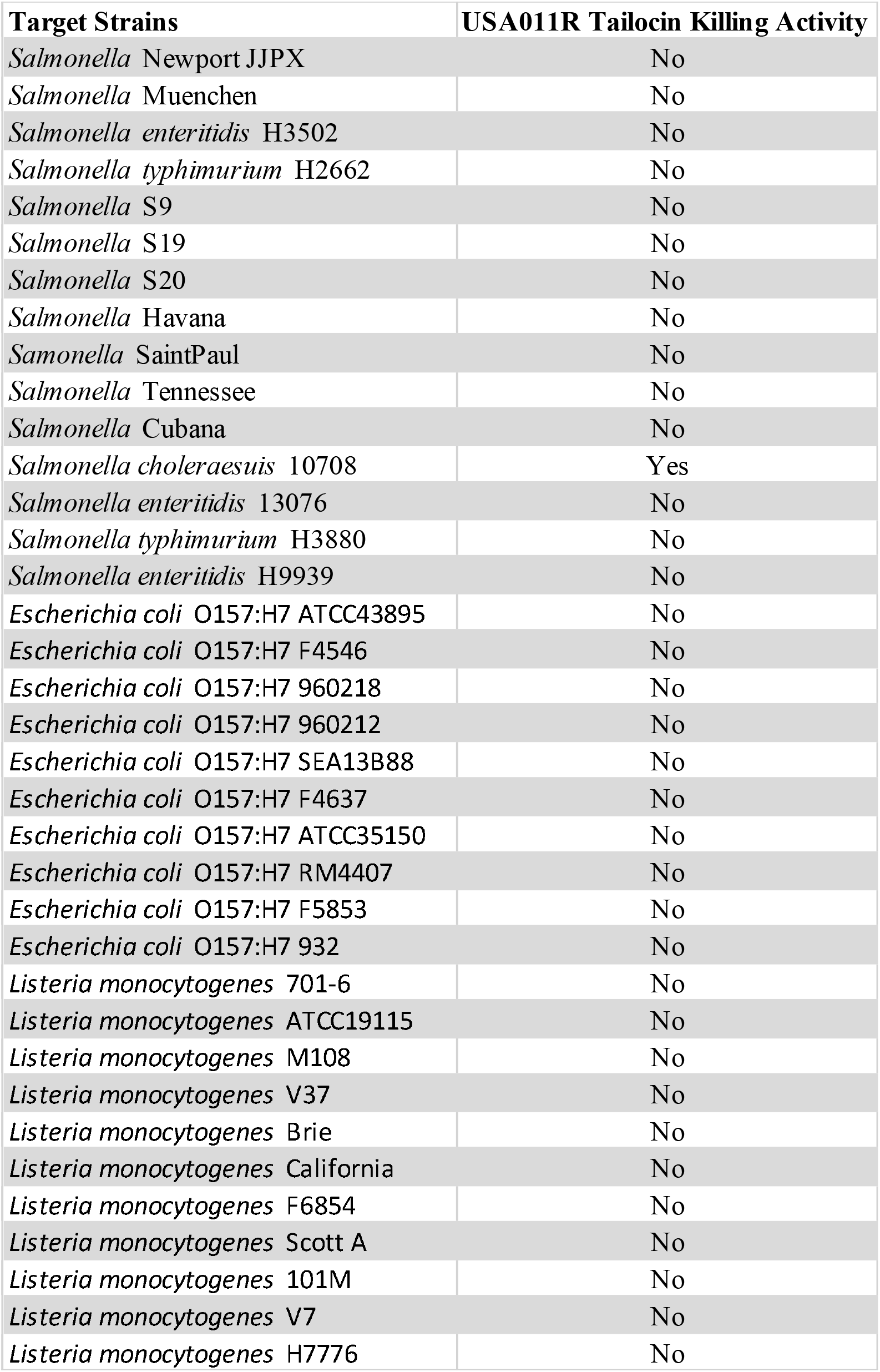
Genus and Species Specific Killing Activity of the Tailocin from *Pseudomonas syringae* strain USA011R.

### Tailocins from Related Strains Differ from USA011R in Cross-Species Host Range and Efficiency of Killing

To investigate phenotypic conservation and divergence of interspecific tailocin killing activity, an array of tailocins from four closely related strains were tested against the same target strains as shown in Figure 1. As a way to quantify the killing activity of these tailocins, we performed a titer assay utilizing the soft agar overlay technique and by performing serial dilutions of the tailocins against the target strains (Fig. 2) on the same overlay plate. As previously reported, *P. syringae* strains USA011R, TLP2, *Pto*2170, and CC457 possess identical killing spectra against all *P. syringae* strains tested thus far, such as *P. syringae* pv. *syringae* B728a (*Psy*B728a) (Baltrus et al. 2018) and we further confirm killing of all these strains against *Psy*B728a here (Fig. 2). Furthermore, all strains also appear to have the same efficiency of activity against *P. syringae*, as similar levels of killing are seen across the dilutions of each strain. In contrast, although there was visible killing by all strains against *Xanthomonas*, there were noticeable differences in effectiveness against *Erwinia* and *Salmonella*. Specifically, while the tailocin from strain TLP2 maintains similar killing activity as USA011R against *E. amylovora*, there is no observable killing of *Erwinia* by tailocins from either *Pto*2170 or CC457. Furthermore, only the tailocin from strain USA011R possesses killing activity against *S. choleraesuis*.

**Figure 2:**
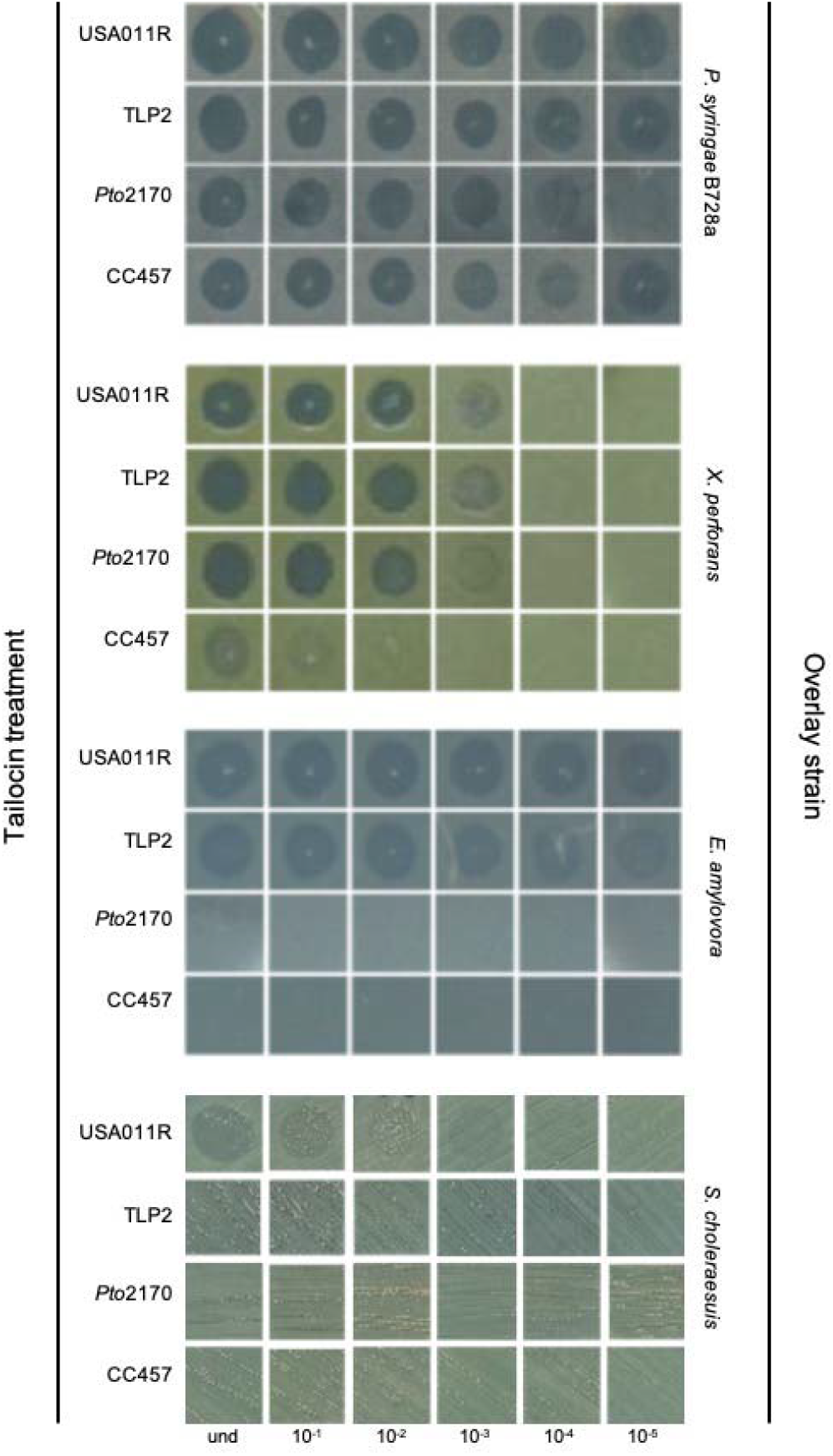
Tailocins from Closely Related Strains Diverge in Interspecies Killing Activity. Titer assays were carried out using the overlay technique as described in Hockett and Baltrus (2017) and modified by performing serial dilutions of tailocins. Tailocins from *P*. syringae strains USA011R, CC457, TLP2, and *Pto*2170 were diluted 10-fold out to 10^−6^ and shown on the horizontal axis. The same tailocin preparations from each strain were used in assays against each of the target strains, with tailocin assays against each strain carried out on the same plate for all tailocin preparations shown. Data is representative of at least three independent overlay experiments. Pictures used for this figure were modified to enhance contrast, but original pictures can be found as supplemental figures S1A, S1B, S1C, and S1D.

Since the same tailocin preparations and dilutions were used for all comparisons shown in Figure 2, and since these tailocins were all plated against the same lawn of target cells, evaluation of tailocin killing by plating dilutions of tailocins also allows for relative assessment of tailocin killing efficiency across all strains. Whereas all tailocins maintain extensive killing activity through the 10^−6^ dilution against *P. syringae*, differentiation of the efficiency of tailocin killing differs across strains against *Xanthomonas*. Although tailocin activity is clearly seen through the 10^−4^ dilutions for USA011R, TLP2, and *Pto*2170, tailocin activity from strain CC457 is only clearly seen to the 10^−2^ dilution. Comparison of this data therefore suggests that the tailocin from CC457 is as effective as those from the other strains against *P. syringae*, but at least ∼10 to ∼100-fold less effective than those of either USA011R, TLP2, or *Pto*2170 against *Xanthomonas*. Put differently, it appears to take 10 to 100-fold more tailocin molecules of CC457 to kill the same amount of *Xanthomonas* cells as the other three strains even though these tailocins appear equally as efficient in killing *P. syringae*.

### Receptor Binding Protein and Chaperone Sequences Show Allelic Diversity Across Strains

Since tailocin killing activity is abolished when both Rbp and chaperone genes are deleted from the tailocin operon in USA011R, we investigated allelic diversity of the protein sequences of these loci across all strains shown in Figure 2. The Rbp protein alignments of USA011R vs. TLP2 and CC457 vs. TLP2 show high similarity in protein sequences with sequence identities of 97% (14 amino acid changes) and 95% (26 amino acid changes), respectively. In contrast, the Rbp and chaperone of *Pto*2170 are highly diverged from those of the other three strains and are also significantly longer (Rbp) and shorter (chaperone) from those in the other three strains. The sequence identity between USA011R and *Pto*2170 is only about 45% (275 amino acid changes), but it is important to take into account that *Pto*2170 contains an addition of 62 amino acids in the Rbp compared to USA011R (Fig. 3A). The chaperone protein alignments of USA011R vs. TLP2 and CC457 vs. TLP2 again show little allelic diversity with sequence identities of 96% (7 amino acid changes) and 89% (21 amino acid changes), respectively. The *Pto2170* chaperone contains 60 amino acids total compared to 189 amino acids of the USA011R chaperone and therefore only shares about 7% sequence identity (Fig. 3B).

**Figure 3:**
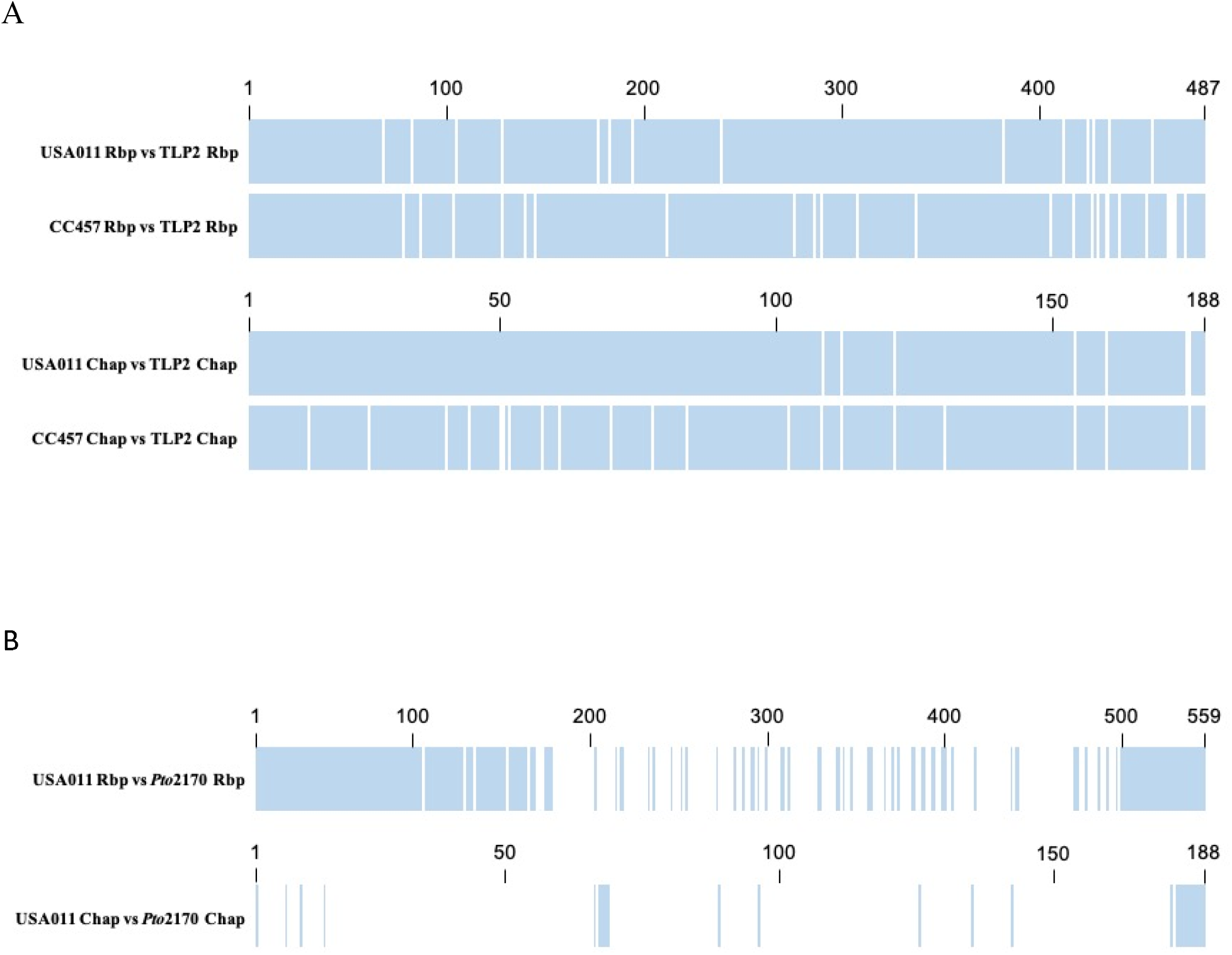
Allelic Diversity in Receptor Binding and Chaperone Proteins Across Closely Related *Pseudomonas syringae* Strains. The Rbp and chaperone protein sequences were extracted from the genomes of *P. syringae* USA011R, CC457, TLP2, and *Pto*2170 and aligned against each other using Geneious Server™ as shown here in this figure. **A)** Rbp and chaperone protein alignments were carried out comparing strains that show limited allelic diversity; USA011R vs. TLP2 and CC457 vs. TLP2. **B)** Rbp and chaperone protein alignments were carried out comparing strains that show greater allelic diversity; USA011R vs. *Pto*2170. In both **A** and **B**, the blue regions indicate shared amino acids and the white regions indicate amino acid changes. Scale on the horizontal axis indicates amino acid position.

## Discussion

Tailocin killing activity across target strains is widely thought to be highly specific and precise, but a handful of previous studies have shown the killing spectra of *Pseudomonas* tailocins could be broader than previously expected (Príncipe et al. 2018; Blackwell and Law 1981; Yao et al. 2017). Our results demonstrate that cross-species killing activity also exists for a subset of *P. syringae* tailocins against the phytopathogenic species *X. perforans* and *E. amylovora* and the human pathogenic species *S. choleraesuis*. However, we also show that this killing activity is not completely conserved across tailocins produced by phylogenetically closely-related strains. It is important to note that, while our previous studies strongly implicate the Rbp and chaperone as main determinants of host range, and allelic diversity in these molecules mirrors divergence in tailocin activity, experiments to definitively match allelic diversity in these loci with tailocin host range are ongoing. It is also possible that other changes in the Rbp sequence or other regions of the genome could be responsible for the altered killing activity, possibly at the single nucleotide level. In contrast, comparisons of these loci in strain *Pto2170* to these sequences within the other three strains, suggests that more extensive changes could also be implicated in tailocin host range differentiation. Discovering the mutations that are causative of phenotypic changes in killing spectra between strains would help us better understand allelic diversification in tailocin-producing strains and could be beneficial for designing tailocins with customized specificities for killing.

Our current knowledge suggests that tailocins utilize the Rbp to bind to the lipopolysaccharide (LPS) of the target (Hockett et al. 2017). Although a single sugar moiety, such as rhamnose, is implicated in tailocin binding by R-type pyocins, previous research has demonstrated that larger scale context and changes in the LPS also dramatically alter tailocin host range (Kandel, Baltrus, and Hockett 2020; Jayaraman et al. 2020). Each of the target species tested here likely have different O-antigen structures even though they all do contain D-rhamnose in their LPS (Maldonado, Sá-Correia, and Valvano 2016; Jayaraman et al. 2020; Lerouge and Vanderleyden 2002). It is also important to note that the observed inter-specific killing activity is also somewhat specific, as the tailocin from USA011R maintains activity against *S. choleraesuis*, but no other tested foodborne pathogens including *Salmonella, E. coli* O157:H7, or *Listeria monocytogenes* strain (Table 1). Furthermore, it is currently thought that tailocins maintain a “one hit, one kill” efficiency against target strain, but, in the least, our data suggest that the story is more complicated. Although the tailocin preparations from CC457 maintain similar efficiency of killing as those from USA011R against *P. syringae* targets, these same preparations are less effective against *Xanthomonas* when the lawns of bacteria are subjected to the same amounts of tailocins (Fig. 2). This result can be explained by hypothesizing that it takes ∼10 to ∼100 fold higher concentrations of tailocin from CC457 compared to USA011R to kill *Xanthomona*s, potentially because tail fibers from CC457 do not bind as well to the LPS of *Xanthomonas*.

Given our data, an overarching question remains; why does interspecies killing activity exist for tailocins? It is thought that bacteriocins are used to outcompete closely related strains which for *P. syringae* likely includes other phytopathogens that could occupy similar niches *in planta* (Ehau-Taumaunu and Hockett 2022), although the strength of selection for this activity is arguable given the frequency that strains might encounter each other and diversity of imagined niches across these species. It is also certainly possible that LPS conformation is selected based on interactions of these species with plants themselves (Newman et al. 2000; Rapicavoli et al. 2018; Ormeño-Orrillo et al. 2008), and that observed killing is simply a byproduct of selection for other phenotypes which in turn sensitize strains to tailocins. For instance, it is possible that a specific conformation of LPS enables each of these strains to evade plant defenses and that specific LPS changes that are selected to better infect plants uncover sensitive rhamnose moieties. In this scenario, selection for interspecies killing activity of USA011R would not be attributed to competition against these species but rather due to convergence of LPS conformation and a pleiotropic tradeoff with tailocin sensitivity.

Tailocins are often thought to have very precise and specific killing activities against conspecific strains. Here we demonstrate that the killing activity of tailocins from *P. syringae* strain USA011R is broader than previously appreciated, with demonstrated activity against some strains classified as *Erwinia, Xanthomonas*, and *Salmonella*. Furthermore, both the killing efficiency and activity of interspecies tailocin killing diverges are not conserved in closely related strains, which hints at additional complexity in tailocin-target interactions. A next step will be to further dissect the molecular basis of differences in interspecific killing activity, with the hopes of potentially developing specific and programmable prophylactic antimicrobials that can be used in agriculture.

## Methods

### Bacterial Strains and Growth Conditions

*P. syringae* strain USA011R was originally isolated by Cindy Morris (Morris et al. 2010). *P. syringae* strain USA011R was derived and isolated in the Baltrus lab by plating out overnight cultures of strain USA011R on King’s B (KB) medium plus rifampicin agar plates and selecting a single colony. The single colony was then grown overnight in KB media plus rifampicin and frozen in 40% glycerol at -80°C. The creation of DBL1424 (a Rbp/chaperone deletion mutant of USA011R) and DBL1701 (a complementation strain derived from DBL1424) was described in (Baltrus et al. 2022). These two genes are located at the 3’ end of a predicted tailocin operon and have also been shown that they are oftentimes exchanged together, allowing for simplicity of complementation (Baltrus et al. 2018). *Xanthomonas perforans* strain DBL942 was isolated by the Baltrus lab from diseased pepper plants in 2014 (DA Baltrus, unpublished). A single colony was grown overnight in Luria-Bertani broth and frozen in 40% glycerol at -80°C. *Erwinia amylovora* ATCC 29780 was obtained from Dr. Julianne Grose at Brigham Young University. All *Salmonella, Escherichia coli* O157:H7, and *Listeria monocytogenes* strains were obtained from the Ravishankar lab at The University of Arizona.

### Bacteriocin Induction and Production

The basic soft agar overlay assay has been previously described in detail (Hockett and Baltrus 2017). For the induction and isolation of R-type syringacins, a single colony of the strain of interest is picked, transferred to 2-3 mL of KB (King’s B medium), and grown overnight on a shaking incubator at 220 rpm and 27°C. The next day, the culture of the strain of interest is back-diluted 1:100 in 3 mL of KB and placed back on the shaking incubator for 3-4 hours. After this incubation period, mitomycin C is added to the 3 mL culture at a concentration of 0.5 µg/mL and incubated on the shaker overnight to allow for induction of bacteriocin production. The next day, 1-2 mL of the mitomycin C induced cultures is centrifuged at 20,000 x g for 5 minutes to form a pellet. The supernatant is carefully transferred to a sterile microcentrifuge tube and 100 µL of chloroform is added per 1 mL of supernatant for sterilization. Immediately after adding chloroform, the sample is vortexed for 15 seconds and then incubated at room temperature for 1 hour. After incubation, the upper aqueous layer is transferred to a new, sterile microcentrifuge tube and is left uncapped at room temperature for several (3-4) hours in the fume hood to allow any residual chloroform to evaporate. Supernatants can be stored at 4°C.

### PEG Precipitation

Polyethylene glycol (PEG) precipitation is used for the separation and concentration of high molecular weight R-syringacin compounds. Final concentrations of 1 M NaCl and 10% PEG are added to the sterile supernatant, inverted repeatedly until dissolved, and incubated for either 1 hour on ice or overnight at 4°C. The samples are then centrifuged at 16,000 x g for 30 minutes at 4°C. After decanting the supernatant, the pellet is resuspended in a volume of 1/10 of the original supernatant volume of buffer (10 mM Tris, 10 mM MgSO_4_, pH 7.0). To remove any residual PEG, an equal volume of chloroform is added to the resuspended pellet, vortexed for 10-15 seconds, centrifuged for 20,000 x g for 5 minutes, and the upper aqueous layer is transferred to a clean microcentrifuge tube. This is repeated for a total of two extractions until no white interface is seen between the aqueous and organic phase layers. Any residual chloroform is allowed to evaporate in a fume hood by leaving the tube uncapped for several (3-4) hours.

### Soft Agar Overlay

A single colony of the target strain is picked, transferred to 3 mL of KB liquid, and grown overnight on a shaking incubator at 220 rpm and 27°C. The next day, the target strain culture is back-diluted 1:100 in 3 mL of KB and incubated for 3-4 hours. Sterile 0.4% water agar is melted and 3 mL is transferred to a clean culture tube and allowed to cool until it is warm to touch. An aliquot (100 µL) of the target strain culture is transferred to the melted water agar and vortexed for a few seconds to thoroughly mix. The water agar mixture is then poured onto a solid KB agar plate, swirled to cover the entire plate, and then allowed to cool for 15-20 min. Once solidified, 5 µL of the PEG-prepared supernatant is spotted onto the dish and allowed to dry. For the titer overlay assay, a serial dilution of each PEG-prepared tailocin supernatant is carried out to 10^−5^ dilution and 5 µL of each dilution is spotted onto the soft agar layer and allowed to dry. The plate is allowed to grow for 1-2 days and then the presence or absence of killing activity is recorded. Soft overlay assays are repeated twice using independent bacteriocin inductions and overlays.

